# Model-based cell clustering and population tracking for time-series flow cytometry data

**DOI:** 10.1101/690081

**Authors:** Kodai Minoura, Ko Abe, Yuka Maeda, Hiroyoshi Nishikawa, Teppei Shimamura

**Author notes:** Equal contributor (Kodai Minoura, Ko Abe).

## Abstract

**Motivation:** Modern flow cytometry technology has enabled the simultaneous analysis of multiple cell markers at the single-cell level, and it is widely used in a broad field of research. The detection of cell populations in flow cytometry data has long been dependent on “manual gating” by visual inspection. Recently, numerous software have been developed for automatic, computationally guided detection of cell populations; however, they are not designed for time-series flow cytometry data. Time-series flow cytometry data are indispensable for investigating the dynamics of cell populations that could not be elucidated by static time-point analysis.

Therefore, there is a great need for tools to systematically analyze time-series flow cytometry data.

**Results:** We propose a simple and efficient statistical framework, named CYBERTRACK (CYtometry-Based Estimation and Reasoning for TRACKing cell populations), to perform clustering and cell population tracking for time-series flow cytometry data. CYBERTRACK assumes that flow cytometry data are generated from a multivariate Gaussian mixture distribution with its mixture proportion at the current time dependent on that at a previous timepoint. Using simulation data, we evaluate the performance of CYBERTRACK when estimating parameters for a multivariate Gaussian mixture distribution, tracking time-dependent transitions of mixture proportions, and detecting change-points in the overall mixture proportion. The CYBERTRACK performance is validated using two real flow cytometry datasets, which demonstrate that the population dynamics detected by CYBERTRACK are consistent with our prior knowledge of lymphocyte behavior.

**Conclusions:** Our results indicate that CYBERTRACK offers better understandings of time-dependent cell population dynamics to cytometry users by systematically analyzing time-series flow cytometry data.

## Background

Flow cytometry is a widely used technology for identifying and quantifying cellular properties and cell populations by measuring expression levels of surface and intracellular proteins at the single-cell level. Modern flow cytometers allow the simultaneous detection of nearly 20 protein markers per cell with a throughput of thousands of cells per second. The flow cytometry technique has greatly contributed to understanding the cellular biological processes and supporting clinical diagnoses in fields including immunology, cancer biology, and regenerative medicine [1, 2, 3].

An important challenge in the analysis of flow cytometry data is the classification of individual cells into canonical cell types, that is, subset populations such as T and B cells. The traditional approach of “manual gating” is performed by visually inspecting a two-dimensional scatter plot, but it suffers from several major limitations, including subjectivity, operator bias, difficulties in detecting unknown cell populations, and difficulties in reproducibility [4, 5, 6].

To overcome these limitations, several methods have been proposed for the computationally guided or automated detection of unknown cell populations by unsupervised clustering, including FlowSOM, X-shift, PhenoGraph, Rclusterpp, and flowMeans [7]. Although these methods have been successfully applied to identify both major and rare cell populations, they are not designed for modeling and analyzing time-series data and thus cannot capture the time-dependent properties and dynamics of cell populations. For example, in clinical applications such as cancer immunotherapies, we are interested in investigating drug effects on cell populations by monitoring their dynamics throughout the treatment period [8]. Time-series flow cytometry data offer information on longitudinal cell population dynamics that could not be elucidated by conventional static time-point data. However, such research is currently limited by a lack of a systematic mathematical framework to adequately model and analyze time-series flow cytometry data.

To address this problem, we propose a new statistical framework, named CYBERTRACK (CYtometry-Based Estimation and Reasoning for TRACKing cell populations), for the automatic clustering and tracking of a mixture proportion of cell populations in time-series flow cytometry data. Our contributions are summarized as follows:

- Our framework is based on the Topic Tracking Model proposed by Iwata *et al.,* 2009, which is designed for tracking topic distribution that changes over time. We extend their model to handle time-series flow cytometry data, which is assumed to follow a multivariate Gaussian mixture distribution.
- By assuming that the mixture proportion at the current time is dependent on that at a previous time, CYBERTRACK is capable of estimating the longitudinal transition of multiple cell populations and detecting the “change-point” in the overall mixture proportion.
- We provide a simple and efficient learning procedure for the proposed model by using a stochastic EM algorithm, which is an alternate iteration of Gibbs sampling and maximum a posteriori (MAP) estimation of parameters. CYBERTRACK is implemented in an R environment, and the implementation is available from https://github.com/kodaim1115/CYBERTRACK.

A conceptual view of an analysis by CYBERTRACK is shown in Figure 1.

**Figure 1.**
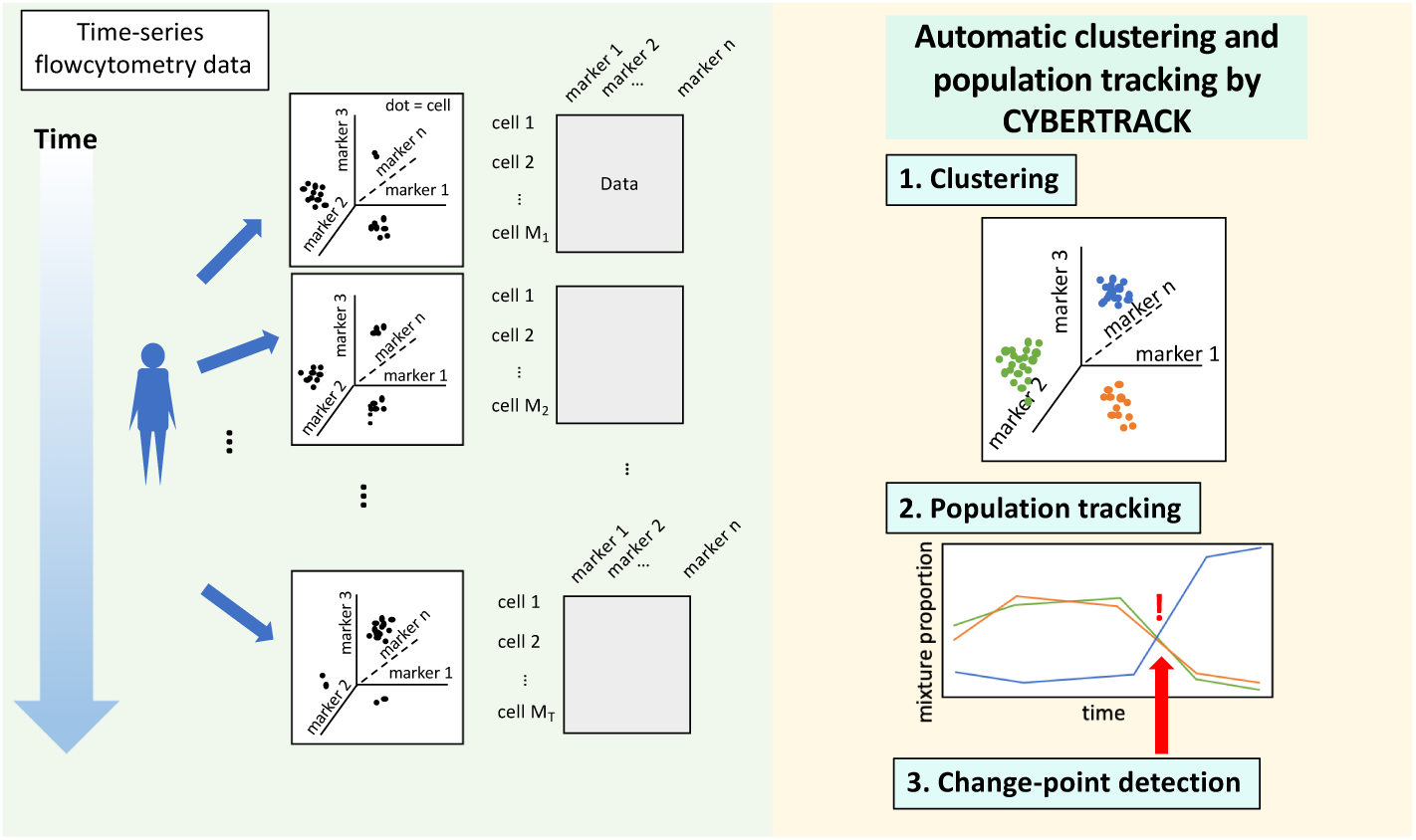
Conceptual view of the analysis by CYBERTRACK. The aim of CYBERTRACK is to model and analyze time-series flowcytometry data to understand dynamic cell population behavior that spans certain period of time. In time-series flow cytometry analysis, cells are acquired sequentially and their expression levels of marker proteins are analyzed, giving data matrices of cells and markers for each timepoint. CYBERTRACK takes these data matrices as an input and performs *i*) clustering, *ii*) tracking cell population dynamics, and *iii*) detecting change-points in cell population constitution.

Our model and algorithm are described in the “Methods” section. To validate its performance and practicability, we applied CYBERTRACK to both simulation and real time-series flow cytometry datasets for two immunological experiments.

## Methods

### Model

Suppose that we observe time series flow cytometry data *y*_*t,d,n*_ ∈ ℝ^*K*^, where *t* ∈ {1, *…, T*} is a time index, *d* ∈ {1, *…, D*} is a case index, *n* ∈ {1, *…, N*_*t,d*_} is a sample index, and *K* is the number of markers. Here, *N*_*t,d*_ represents the number of samples observed at time *t* for case *d*. The objective of this study is to perform clustering of samples and track the time-dependent transition of cluster mixture proportion *Π*_*t,d,l*_ for each case, where *l* ∈ {1, *…, L*} is a cluster index. Our model is inspired by the Topic Tracking Model, and is an extension of the multivariate Gaussian mixture model. Topic Model is a Bayesian model which that was originally designed to extract latent semantics, or “topics”, from text data. Topic Tracking Model is an extension of Topic Model specialized in tracking time-varying topic distribution [9]. Although the original Topic Tracking Model assumed each word was generated from a multinomial distribution, this assumption does not apply to the case with flow cytometry data. Therefore, we assumed that flow cytometry data follow a multivariate Gaussian mixture distribution, and we constructed the algorithm to estimate parameters. Here, topics correspond to cell populations such as T cells or B cells. Figure 2 illustrates a plate diagram of our proposed model, where, *z*_*t,d,n*_ is a latent cluster vector of length *L* that holds 1 for the *l*-th element when a sample is generated from cluster *l* and holds 0 otherwise. We assume that each sample is generated from a multivariate Gaussian mixture distribution with the parameter vector ***μ***_*l*_ and **Σ**_*l*_, which represents the mean and the covariance matrix for cluster *l*, respectively. More specifically, the generative process of CYBERTRACK is defined by

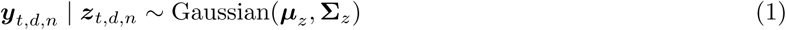

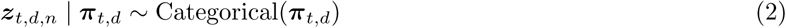

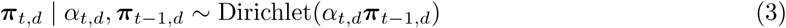

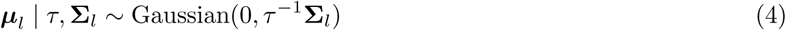

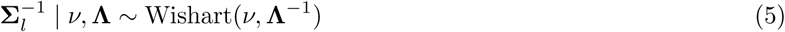

where *z* is a latent cluster of the *n*-th sample at time *t* for case *d* indicated by *z*_*t,d,n*_, ***μ****z* and **Σ**_*z*_ are the mean vector and covariance matrix of the latent cluster, respectively, 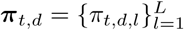 is the mixture proportion vector, and *α*_*t,d*_ represents the persistency parameter, which indicates how consistent the mixture proportion at time *t* is compared with that at the previous time *t -* 1. A smaller *α*_*t,d*_ value indicates a larger discrepancy between the mixture proportion at time *t* and *t -* 1. Thus, timepoints with relatively small persistency parameters could be considered as “change-points” in the mixture proportion. 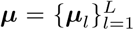 is the mean vectors of clusters, and 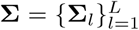 is the covariance matrices of clusters. *τ* is the hyperparameter of the ***μ*** prior distribution, and **Λ** and *v* are the hyperparameters of the **Σ** prior distribution.

**Figure 2.**
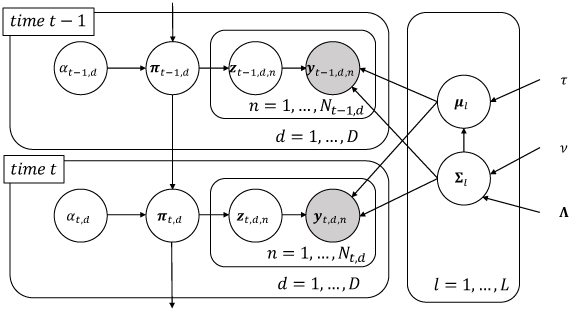
Graphical model of CYBERTRACK.

### Parameter Estimation

Parameter estimation in CYBERTRACK is based on the stochastic EM algorithm, which is an alternate iteration of Gibbs sampling and maximum a posteriori estimation of parameters. Suppose *t* is the current time, and suppose we have flow cytometry data matrix 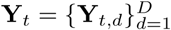 and a mixture proportion matrix 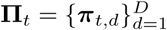, where 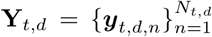. We perform the inference of latent clusters based on Gibbs sampling. Let 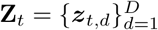 be the set of latent clusters of all cases at time *t*, where 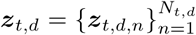. The posterior distribution of **Z**_*t*_ given **Y**_*t*_, *Π*_*t*_, ***μ***, and **Σ** can be written as follows:

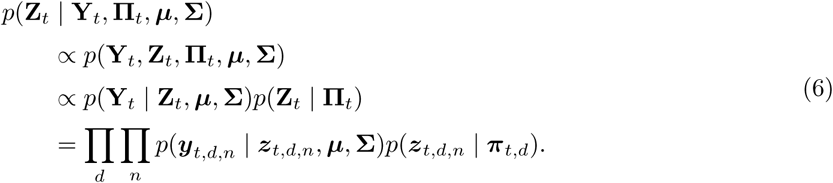

The logarithm of above will be:

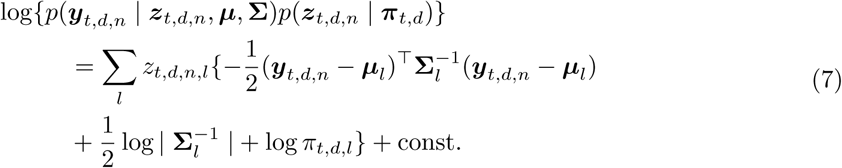

Therefore, ***z***_*t,d,n*_ is sampled from the following categorical distribution:

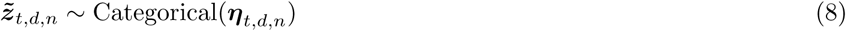

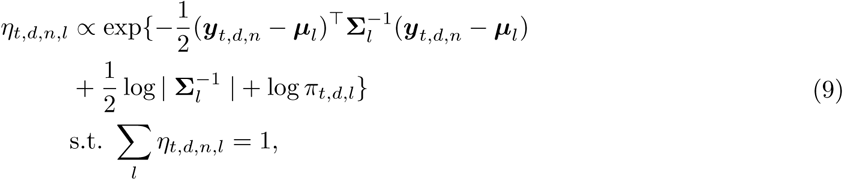

where 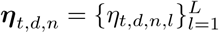. Suppose we have the mean of the previous mixture proportion 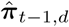. The persistency parameter *α*_*t,d*_ is estimated by fixed point iteration.

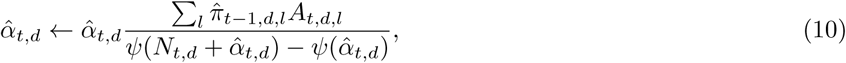

where 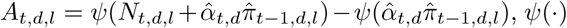 is the digamma function 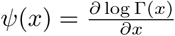, and *N*_*t,d,l*_ is the number of samples assigned to cluster *l* at time *t* for case *d*. The mean of π_*t,d,l*_ is then calculated as follows:

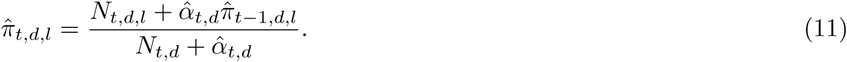

We substitute the E-step of the EM algorithm by Gibbs sampling, then ***μ*** and **Σ** are updated in the M-step is as follows:

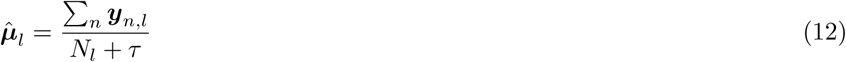

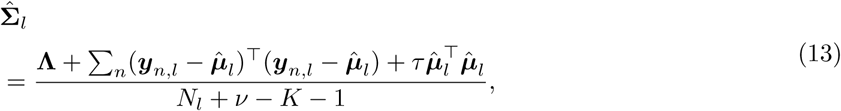

where ***y***_*n,l*_ is the *n*-th sample assigned to cluster *l*, and *N*_*l*_ is the number of samples assigned to cluster *l*.

## Result

### Simulation Study

We conducted a simulation experiment to examine the performance of CYBERTRACK. We set *K* = 10, *T* = 5, and *D* = 2. The ***µ*** and **Σ** was randomly generated. **π** were set manually to have change-points for mixture proportions. For the hyperparameters, we set *τ* = 10^-5^ and *ν* = *K* + 2, and **Λ** was set to an identity matrix, which is equivalent to giving weakly informative priors. With this parameter setting, we randomly generated 1000 samples for each timepoint in each case (10,000 samples in total). The simulation was repeated 10 times, and different data was synthesized each time. The mean and standard error (se) for the estimated 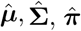, and 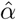 are shown in Figure 3. The results show that the parameters for a multivariate Gaussian mixture distribution were reasonably estimated by the stochastic EM algorithm, and that CYBERTRACK successfully tracked the time-dependent transition of the mixture proportion in multiple cases. As shown in Figure 3d, 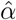 holds small values at *t* = 3, 5 for case 1 and *t* = 2, 4 for case 2, indicating the dramatic transition of the mixture proportion at that timepoint.

**Figure 3.**
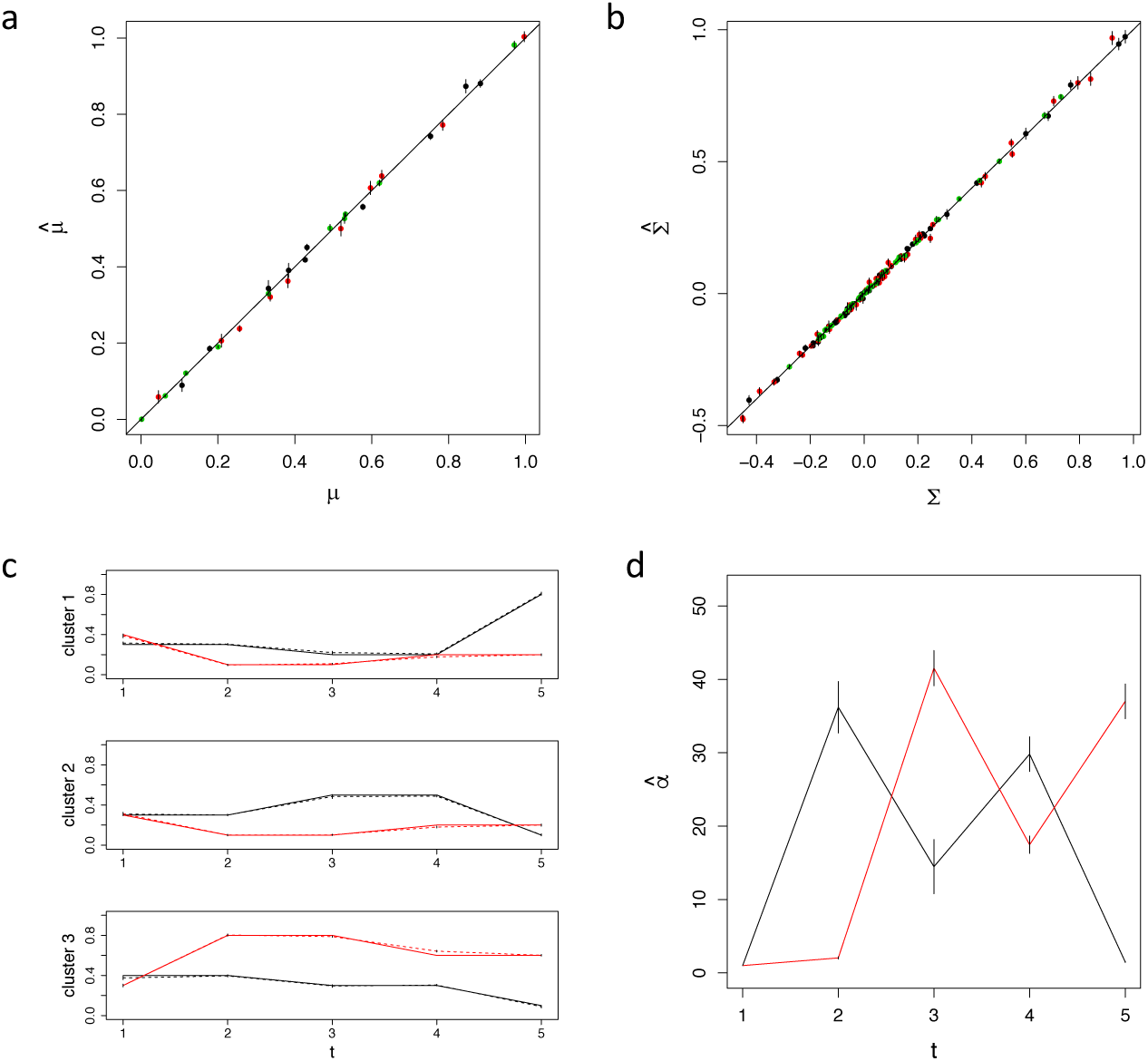
Simulation result for CYBERTRACK analysis. **a**, Estimated 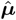 were plotted against true ***µ***. Each dot represents elements of 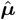 and was color-coded by cluster. **b**, Estimated 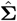 values were plotted against true **Σ**. Each dot represents elements of 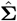 was were color-coded by cluster. **c**, Estimated 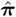 for simulation data. Black and red lines represent the proportion mixture for case 1 and 2, respectively. Solid lines indicate the true mixture proportion and dashed lines indicate estimated proportion. **d**, Black and red lines represent proportion mixtures for cases 1 and 2, respectively. Timepoints where the alpha values decrease substantially indicate change-points in the overall mixture proportion. Error bars represent standard error.

### Results for Real Data

To validate the CYBERTRACK performance for cell clustering and tracking mixture proportions of cell populations, we applied CYBERTRACK to real world flow cytometry data uploaded to Cytobank (https://www.cytobank.org/). In Landrigan’s study (https://community.cytobank.org/cytobank/experiments/35226), naive CD4+ T cells were purified and stimulated using anti-CD3 and anti-CD28 antibodies. Five cases were tested: unstimulated, stimulated by only anti-CD3 antibody, and stimulated by both anti-CD3 and anti-CD28 antibodies, with two dosages tested for the anti-CD3 antibody (0.3 *µ*g/mL and 0.8 *µ*g/mL). It is known that the stimulation of CD3 triggers the activation of naive CD4+ T cells, which accompanies the phosphorylation of SLP76/S6 and CD247 (pSLP76/pS6, pCD246) [10]. CD28 is the co-stimulatory factor that enhances and prolongs T cell activation [11]. Soon after activation, the levels of pSLP/pS6 and pCD247 decrease owing to negative feedback. Consequently, the cells become CD45RO+ memory T cells.

To determine the number of clusters, we used the elbow method, which involves plotting the sum of squared error (SSE) within each cluster against the number of clusters. For Landrigan’s study, the number of clusters was determined as 16 by using the elbow method (Figure 4a). Figure 4b shows the heatmap generated from the estimated 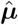 clusters 1, 7, 8, 9, 12, and 16 are the pSLP76/pS6+ pCD247+ activated naive T cells, and clusters 2, 4, 6, and 13 are pSLP76/pS6-pCD247-CD45RO+ memory T cells. The time-dependent transition of the mixture proportion is shown in Figure 5. While the mixture proportion remains stable over time in unstimulated cases, other cases show dynamic fluctuation, as expected. In stimulated cases, a high proportion of activated naive T cells (specifically, cluster 16 and 7 for dosages 0.3 and 0.8 *µ*g/mL, respectively) was observed at *t* = 3 min. Their proportions decreased through *t* = 6, 10 min by the T cell’s negative feedback mechanism. Figure 5 shows that as the number of activated T cells decreases, the memory T cell populations increase, indicating the transformation of naive T cells into memory T cells. This behavior was well represented by 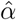 estimates, shown in Figure 6; the 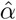 for stimulated cases shows small values at *t* = 6 and *t* = 10 min compared with that of unstimulated cases, indicating dynamic changes in cell population constitution at those timepoints. Interestingly, in cases stimulated with both anti-CD3 and anti-CD28 antibodies, a prominent increase of clusters with moderate levels of pSLP76/pS6 and pCD247 (cluster 14 and 1 for dosages 0.3 *µ*g/mL and 0.8 *µ*g/mL, respectively) was observed at *t* = 6. These clusters can be interpreted as cell populations that are transitioning from a highly activated state to an inactivated memory state. This is consistent with the well-known prolonged T cell activation by stimulation of CD28, thus further indicating that CYBERTRACK is capable of illustrating dynamic biological processes from time-series flow cytometry data [11].

We also applied CYBERTRACK to the data in Huang’s study (https://community.cytobank.org/cytobank/experiments/5002), where cells were collected from mice whose lymph nodes were stimulated with either interleukin 7 (IL7) or interferon alpha (IFN*α*). It is known that IL7 and IFN*α* interact with their receptors on the lyphocytes’ surface and activate lymphocytes through the phosphorylation of STAT family proteins (e.g., pSTAT1 and pSTAT5), which promotes the transcription of immune-related genes [12, 13].

**Figure 4.**
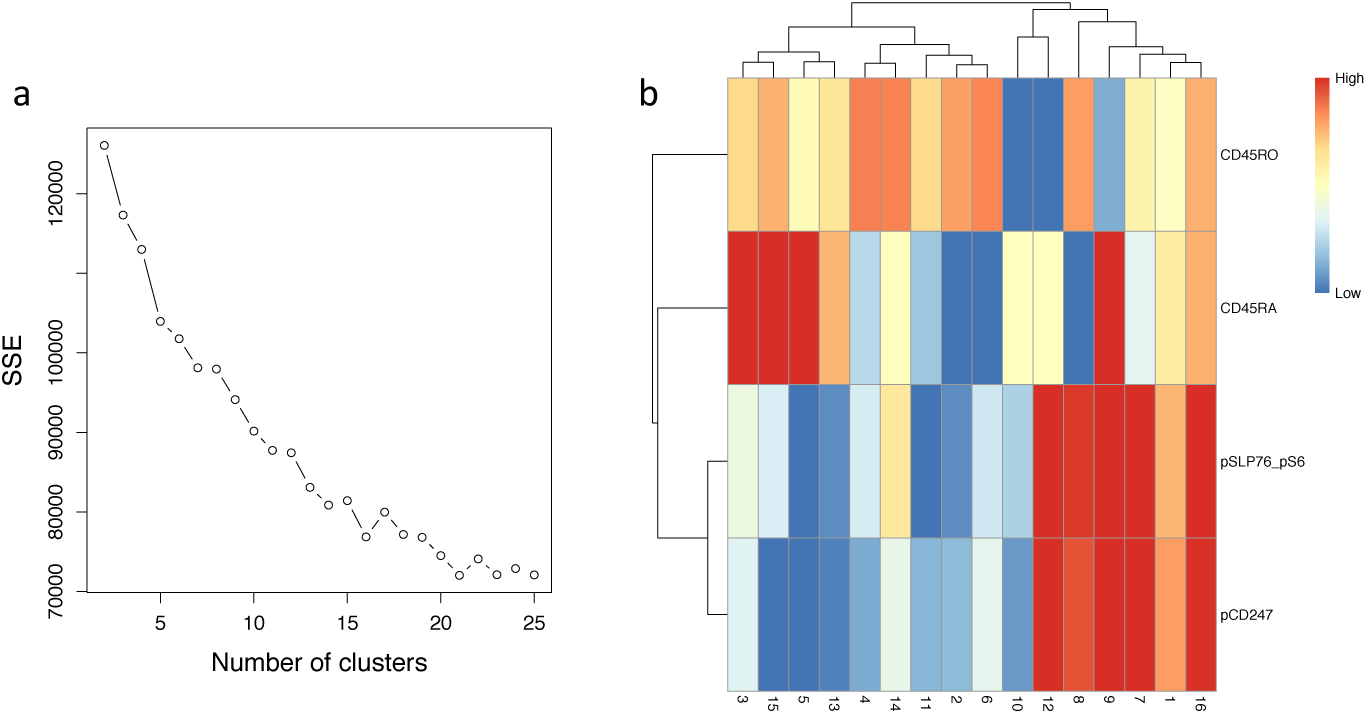
Clustering result for CYBERTRACK analysis on Landrigan’s study. **a**, Elbow plot for Landrigan’s study. SSE saturated at 16 clusters. **b**, Heatmap for Landrigan’s study. Cluster number is shown on the *x* axis and markers are shown on the *y* axis.

**Figure 5.**
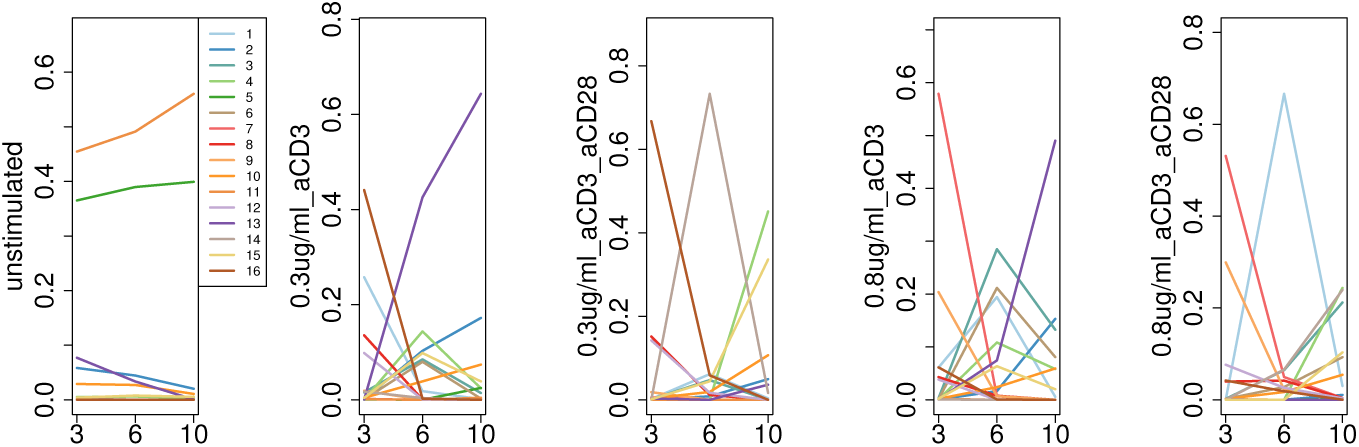
Cell population tracking result for Landrigan’s study. Mixture proportions for five cases were drawn in different graphs. Each colored line represents the mixture proportion for a different cluster. *t* is on the *x* axis and the mixture proportion is on the *y* axis.

**Figure 6.**
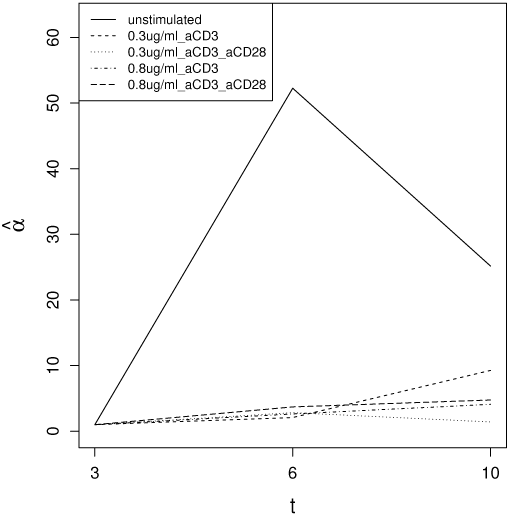
Estimated 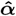 for Landrigan’s study.

The number of clusters was determined as 26 by using the elbow method (Figure 7a), and the heatmap is shown in Figure 7b. In Huang’s study, T cells were identified by CD4 and/or TCR*β*, and B cells were identified by B220. As shown in the heatmap, CYBERTRACK clustered cells into canonical cell types, which include CD4+ TCR*β*+ T cells (clusters 4, 6, 9, 10, 17, and 26), CD4-TCR*β*+ T cells (clusters 2, 3, 15, 24), and B220+ B cells (clusters 1, 3, 5, and 21). Clusters with extremely high levels of both B220 and TCR*β* are thought to be debris; therefore, they were excluded from further interpretation. Figure 8 shows the time-dependent transition of the mixture proportion for each cell population. CYBERTRACK detected cell populations that increased over time in both cases. These cell populations include pSTAT1+ pSTAT5+ T cell (cluster 3) and pSTAT1+ B cell (cluster 5), which are typical cell populations that are known to emerge upon IL7 and IFN*α* stimulation. Furthermore, CYBERTRACK also illustrated cell population dynamics that differed in two cases; pSTAT5+ T cells (clusters 9 and 24) increased only when stimulated by IL7, whereas pSTAT1+ T cells (clusters 6 and 15) increased only in the IFN*α*-stimulated case. Although IL7 and IFN*α* are known to induce the phosphorylation of a variety of STAT family proteins, the result shown here may reflect the preferential upregulation of STAT5 and STAT1 by IL7 and IFN*α*, respectively [14, 15]. The estimated 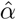 shows that the change-points are located at *t* = 2, 4 min for IL7 stimulation and *t* = 2 min for IFN*α* stimulation (Figure 9). Furthermore, analysis by CYBERTRACK revealed that stimulation by IL7 induces more dramatic changes in cell population constitution at an early stage (until *t* = 4 min), as indicated by the small 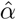 values.

**Figure 7.**
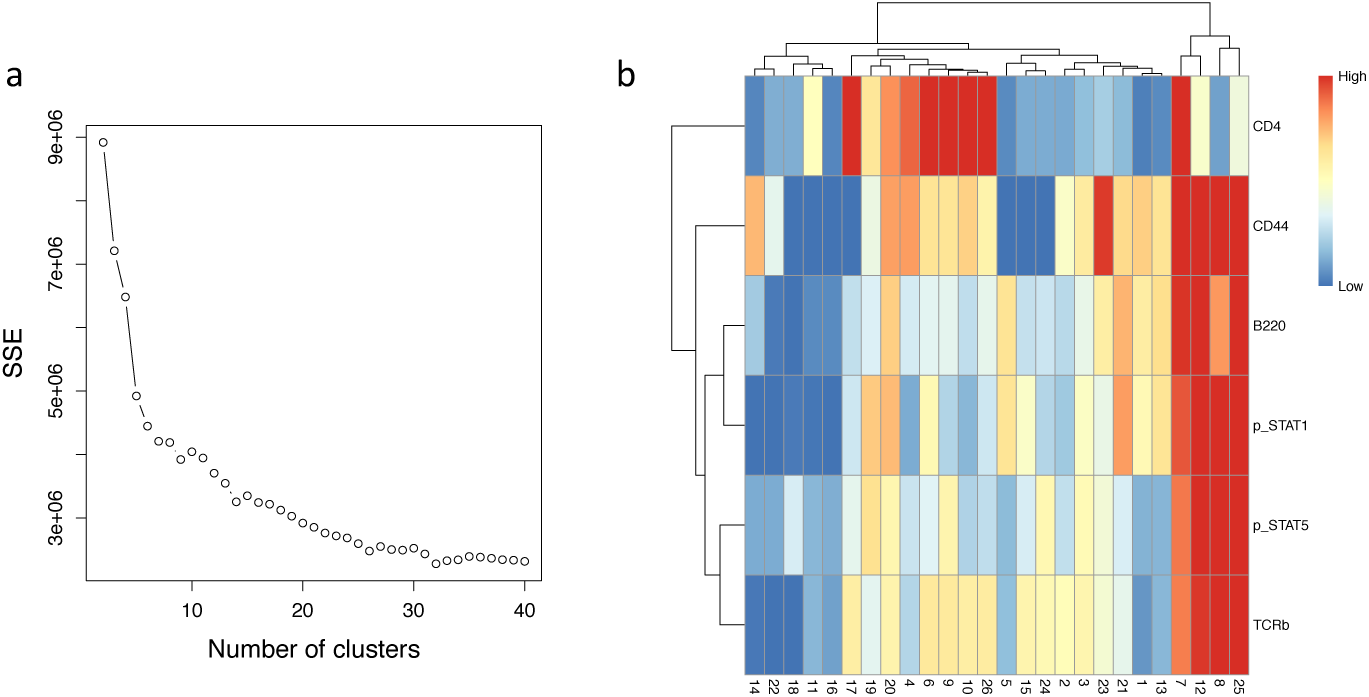
Clustering result for Huang’s study. **a**, Elbow plot for Landrigan’s study. SSE saturated at 16 clusters. **b**, Heatmap for Landrigan’s study. Cluster number is shown on the *x* axis and markers are shown on the *y* axis.

**Figure 8.**
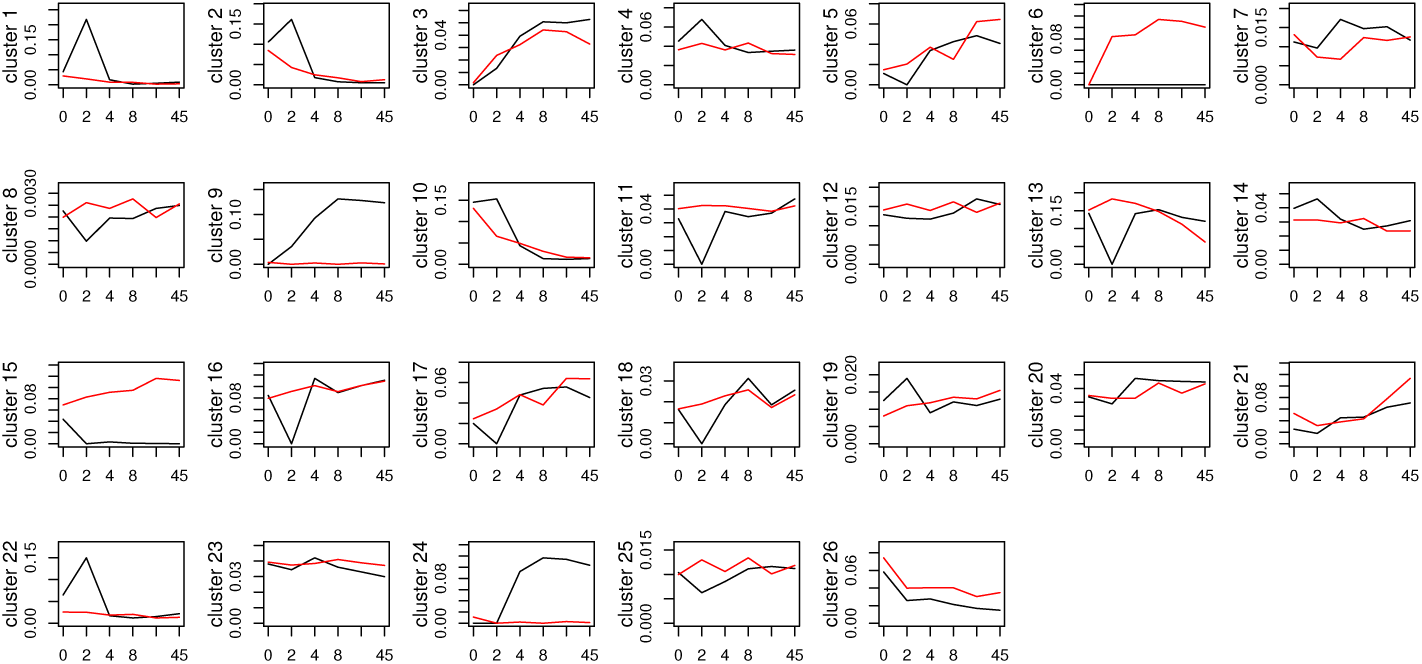
Cell population tracking result for Huang’s study. Mixture proportions for 26 clusters were drawn in different graphs. Black and red lines represent IL7 stimulation and IFN*α* stimulation, respectively. *t* is on the *x* axis and the mixture proportion is on the *y* axis.

**Figure 9.**
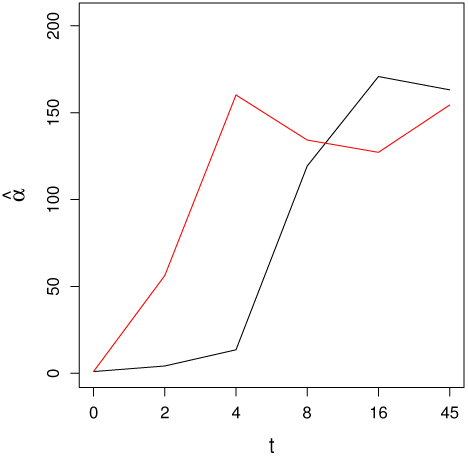
Estimated 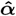 for Huang’s study. Black and red lines represent IL7 stimulation and IFN*α* stimulation, respectively.

## Discussion

Here, we propose a model-based cell clustering and population-tracking algorithm called CYBERTRACK. The aim of CYBERTRACK is to discover the underlying dynamics of cell populations in time-series flow cytometry data. Our model is inspired by the Topic Tracking Model [9], and we modified it for the parameter estimation of a multivariate Gaussian mixture distribution. CYBERTRACK is capable of (*i*) cell clustering, (*ii*) tracking the mixture proportion of each cell population, and (*iii*) detecting the change-point in the overall mixture proportion.

Recently, a tool called mass cytometry was introduced to the field of biomedical research. Mass spectrometry-based detection of marker genes by mass cytometry has enabled the investigation of more than 40 markers simultaneously, providing much more informative data with higher-dimensions compared with fluorescence-based conventional flow cytometry. Recent research trends in single-cell biology highly depend on mass cytometry, and it has contributed to many important discoveries [16]. One limitation of CYBERTRACK is that it is inapplicable to mass cytometry data, because the data generated by mass cytometry do not follow a multivariate Gaussian distribution. Our future aim is to extend CYBERTRACK for application to time-series mass cytometry data.

The application of CYBERTRACK to simulation and real flow cytometry data has validated its performance for cell clustering and tracking mixture proportions in multiple cases. The results of CYBERTRACK analysis using two immunological experiments were consistent with our prior knowledge, which validates CYBERTRACK’s ability to analyze time-series flow cytometry data. We believe that CYBERTRACK will be a powerful tool in various fields involving the investigation of cell population dynamics. For instance, in the field of cancer immunotherapy, the longitudinal immune monitoring of patients has become increasingly important as it provides information on the impact of therapeutic treatment on certain cell populations, or in finding cell populations that can be used as prognostic markers. Furthermore, CYBERTRACK will also be useful in basic research as it can give insights into flow cytometry time-series data in an unbiased manner. Because CYBERTRACK is capable of clustering cells from different cases, it is easy for researchers to compare population dynamics in experiments with a control and several cases.

## Ethics approval and consent to paticipate

Not applicable.

## Consent to publish

Not applicable.

## Availability of data and materials

Our software is available from GitHub (https://github.com/kodaim1115/CYBERTRACK). Data for the immunological experiments used in this paper are available from Cytobank (https://cytobank.org/).

## Author’s contributions

KM, KA, TS designed the proposed algorithm. All authors reviewed and approved the final manuscript.

## Acknowledgements

Not applicable.

